# Remotely sensed indicators and open-access biodiversity data to assess bird diversity patterns in Mediterranean rural landscapes

**DOI:** 10.1101/408187

**Authors:** Inês Ribeiro, Vânia Proença, Pere Serra Ruiz, Jorge Palma, Cristina Domingo, Xavier Pons, Tiago Domingos

## Abstract

Changes in ecosystem area are often used to assess human impacts on habitats and estimate biodiversity change. However, because species respond to structural changes at fine spatial scales the use of area alone may not capture all relevant changes. Operational costs limit the assessment of biodiversity change at a simultaneously fine spatial resolution and large scales. The development of cost-effective and expedite methods to monitor biodiversity change is therefore required. We use open access satellite imagery and biodiversity data to investigate the importance of variables of habitat extent and structure in explaining species richness and community dissimilarity of forest and open-land birds at the regional scale. Moreover, because Mediterranean landscapes are subject to seasonal dynamics, we explore the indicator value of remotely sensed variables measured in spring and summer. A large-scale dataset of bird occurrence data, including 8042 observations and 78 species, distributed by 40 landscape-sized cells, was assembled from GBIF after controlling for data quality. We found that summer satellite imagery, when the green perennial vegetation is more apparent, is particularly suited to model the diversity patterns of forest species, because distribution of tree cover in the landscape is well captured. Summer data is also useful to monitor the perennial elements that shape landscape structure and the habitat of open-land species. Specifically, mean NDVI and a second-order NDVI texture variable, were found to be good indicators of forest and open-land habitats, respectively. The use of spring imagery appears to be useful to monitor habitat structure within open-land habitat patches. Overall, NDVI texture measures were found to be good predictors of bird diversity patterns at large scales. Also, we were able to successfully conduct a regional scale analysis using open-access data, which illustrates their potential to inform large scale biodiversity monitoring.

## Introduction

Species diversity patterns are shaped by multiple factors, including environmental factors, such as climate, primary productivity, habitat area and habitat diversity, and species-specific factors, such as species functional traits and evolutionary history (Rosenzweig, 1995; Desrochers et al., 2011; Martins et al., 2014). The effect of climate and productivity variables is more evident at spatial extents where climatic ranges are broader and variation more pronounced, such as global to regional scales, while the effect of habitat variables gains importance at smaller extents, from regional to local scales (Rosenzweig, 1995; Field et al., 2009; Desrochers et al., 2011; Martins et al., 2014). Regarding species-specific factors, species with different ecological requirements will show different responses to environmental factors and to environmental changes. For instance, the use of species groups based on habitat preferences may help unravel different responses to habitat change within species communities (Proença & Pereira, 2013; Martins et al., 2014; Pereira et al., 2014).

Changes in ecosystem area, documented by airborne and satellite imagery, and by the derived map products, have been used to assess human direct impacts on habitat availability and estimate biodiversity change (Plieninger, 2006; Fischer & Lindenmayer, 2007; Pereira et al., 2010). However, the use of area alone, as an indicator of habitat change may only partially capture changes in habitat availability for species (Hortal et al., 2006; Kallimanis et al., 2008). That is the case when changes at fine spatial grains affect ecosystem structure, but not the main features that define the ecosystem type neither its overall extent. For instance, changes in tree density or vegetation structure, may affect habitat availability, including its quality and structural diversity, within apparently stable forest patches, and consequently affect species presence and diversity patterns (Jankowski et al., 2013; Pereira et al., 2014; Lengyel et al., 2016; Zellweger et al., 2017). Moreover, habitat structure is often described by coarse grain metrics of landscape configuration based on the size, shape, and type of land cover patches, thus overlooking variation at finer spatial grains. Mapping and monitoring ecosystem structure at a fine resolution in large spatial extents poses an operational trade-off, because it requires an intensive sampling effort over a large spatial extent (Wood et al., 2012). Satellite remote sensing offers a tool to overcome such trade-off, as it allows to capture variations in ecosystem structure at fine spatial grains across large spatial extents and with high sampling frequency (Proença et al., 2017; Rocchini et al., 2018). For instance, texture measurements from satellite imagery have been used to measure vegetation and landscape structure (Wood et al., 2012, 2013), and have been found to be a good predictor of species richness patterns (Oindo & Skidmore, 2002; St-Louis et al., 2009, 2014). On the other hand, although texture statistics from optical images inform on vegetation horizontal structure they are not the adequate to capture vertical structure, especially in multilayered systems, notably forests. The inclusion of radar and lidar sensors in satellites, such as the Synthetic Aperture Radar (SAR) and the Light Detection and Ranging (LiDAR), is a major step to address this challenge and produce enhanced spatial layers of habitat structure (Nagendra et al., 2013; Culbert et al., 2012; Zellweger et al., 2017). However, while promising, current data availability from active sensors is limited, and data processing and interpretation still require advanced technical skills that constrain their use (Nagendra et al., 2013; Pettorelli et al., 2016).

This study investigates the importance of variables of habitat extent and structure in explaining the patterns of bird species richness and community dissimilarity at the regional scale. We use a land cover map and optical satellite imagery as data sources of habitat extent and structure. Moreover, we explore the indicator value of remotely sensed variables measured in different seasons. We selected the Alentejo region (NUTS II) in Portugal (Figure 1) as the study area for its extent, landscape heterogeneity and seasonality. Cropland, oak forest and montados (traditional agro-forestry systems) occupy a large share of the territory, providing key habitats for bird communities (Godinho & Rabaça, 2011; Pereira et al., 2015), and forming landscape mosaics with diverse levels of spatial heterogeneity. This heterogeneity is maintained both by the habitat mix and landscape configuration, at coarser grains, and by the variation in vegetation structure at finer grains. In addition to the spatial heterogeneity, these landscapes are also characterized by strong seasonality, typical of the Mediterranean climate, that results in dynamic landscapes (Costa et al., 2009, 2011). A regional scale dataset of bird species occurrences was compiled for this study using up-to-date data made available by GBIF (Global Biodiversity Information Facility; GBIF.org). After checking for data quality, species data were collected in 40 landscape mosaics (Appendix 1), distributed across the region, and divided in two groups of habitat affinity: forest and open-land habitats.

**Figure 1.**
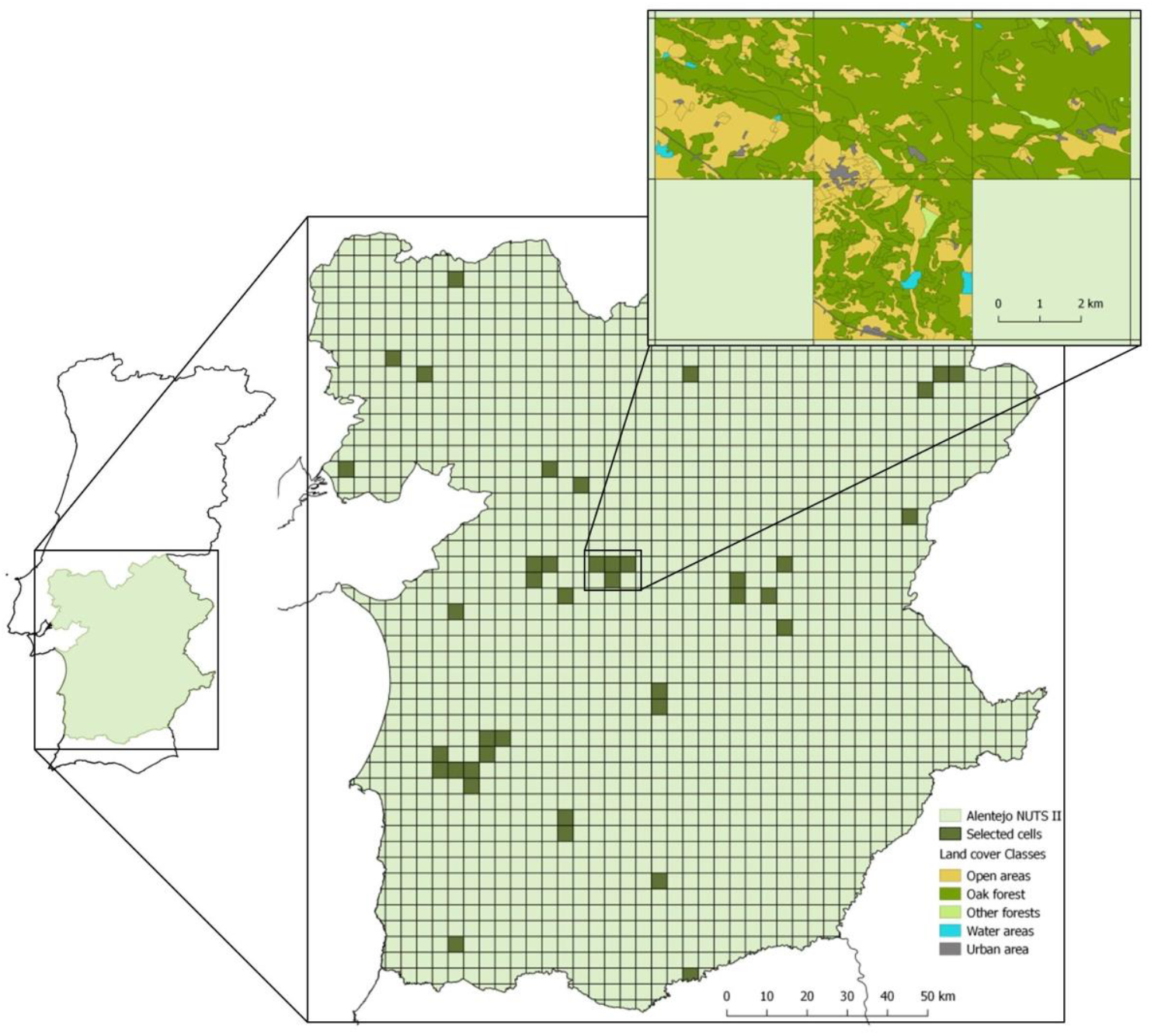
Distribution of the 40 selected cells in Alentejo (i.e., well surveyed cells with eligible land cover, see methods for details on cell selection). An example of the land cover mosaic is shown for four cells.

We expect the richness of a species group in a multi-habitat landscape to be positively related to the extent and structural diversity of their main habitat, with both factors being more important than the overall landscape structure. We also expect habitat extent and structure to be more important than geographic distance or climate when explaining the level of dissimilarity between communities in the region. Hence, we expect habitat descriptors, including remotely sensed variables, to be good indicators of bird diversity patterns at the regional scale. Moreover, we expect summer data, which mostly capture perennial tree cover, to perform better as an indicator of forest species communities, and spring data, which also capture herbaceous cover at its peak productivity, to perform better as an indicator of open-land communities. Finally, open-land species may show stronger responses to habitat structure variables than forest species because these variables were derived from optical remotely sensed data, which do not capture understory vegetation structure.

## Methods

### Study area

Our study area is the Alentejo region (NUTS II) in Portugal (Figure 1). This is a predominantly rural region with low population density. The climate is Mediterranean with hot and dry summers (Kottek et al., 2006). The landscape is mostly plain with extensive land uses, but intensification is increasing with potential impacts on biodiversity (Pinto-Correia & Mascarenhas, 1999; Costa et al., 2011; Plieninger et al., 2015; Gossner et al., 2016). Most notably, montados (dehesas in Spanish), listed under the EU Habitats Directive (92/43/EEC) for their conservation value, cover a significant share of the landscape. These are traditional systems with a silvo-pastoral use, where cork oak (*Quercus suber*) and holm oak (*Q. rotundifolia*) are the dominant trees, forming pure or mix stands with a tree cover varying between 10% and 30% (Caetano et al., 2010). Montados are managed for multiple productive purposes, the most important being cork extraction, pastures and livestock (Pinto-Correia et al., 2011; Bugalho et al., 2011). In addition, these landscapes provide significant opportunities for ecotourism and for recreational activities (Berrahmouni et al., 2007). The multifunctional use promotes habitat structural diversity, which combined with the large regional extent and the generally low human population density enables the persistence of many species, including endangered species (e.g., Iberian lynx, *L. pardinus*, Adalbert’s Eagle, *Aquila adalberti*,). For the purpose of this study, all stands of the Portuguese land cover map, COS 2007 v2.0 (Caetano et al., 2010), dominated by oaks (≥75%), and with a tree density of at least 10% were designated as oak forest. For data analysis, the study area was divided in to landscape-level cells of approximately 4.89 km × 4.89 km (Figure 1, Appendix 1) using a level 5 *geohash* grid (www.geohash.org).

### Study design

To account for the seasonal variation in vegetation cover and ecosystem primary productivity and for differences in habitat use by forest and open-land species, we assigned a distinct set of candidate predictor variables to each species group, which includes NDVI (Normalized Difference Vegetation Index) texture variables measured either in spring or in summer (Table 1). This approach enables testing the differential response of species groups to remotely sensed data collected (1) in spring when the herbaceous vegetation cover is at peak productivity, thus providing a better signal of open-land habitats, or (2) in summer, when the contrast between the senescent herbaceous cover and the perennial vegetation, namely tree cover, is better captured. Moreover, NDVI measurements were assessed at two scales for each grid cell: the landscape scale (i.e., using all pixels in the grid cell) and the habitat scale (i.e., using only the pixels overlapping the preferred habitat, either forest or open-land habitats, in the Portuguese land cover map layer). With this approach we aim to account for species responses to the overall surrounding landscape and to the preferred habitat. In addition to NDVI measures, all sets of candidate variables included other environmental variables, measured at the landscape scale, namely climatic, topographic and land cover variables (Table 1). From the full set of 70 candidate variables (Appendix 2), we retained a final set of non-colinear variables to be used in the analysis of species richness and community dissimilarity patterns. Variable selection is described in the section *Data analysis.*

**Table 1.**
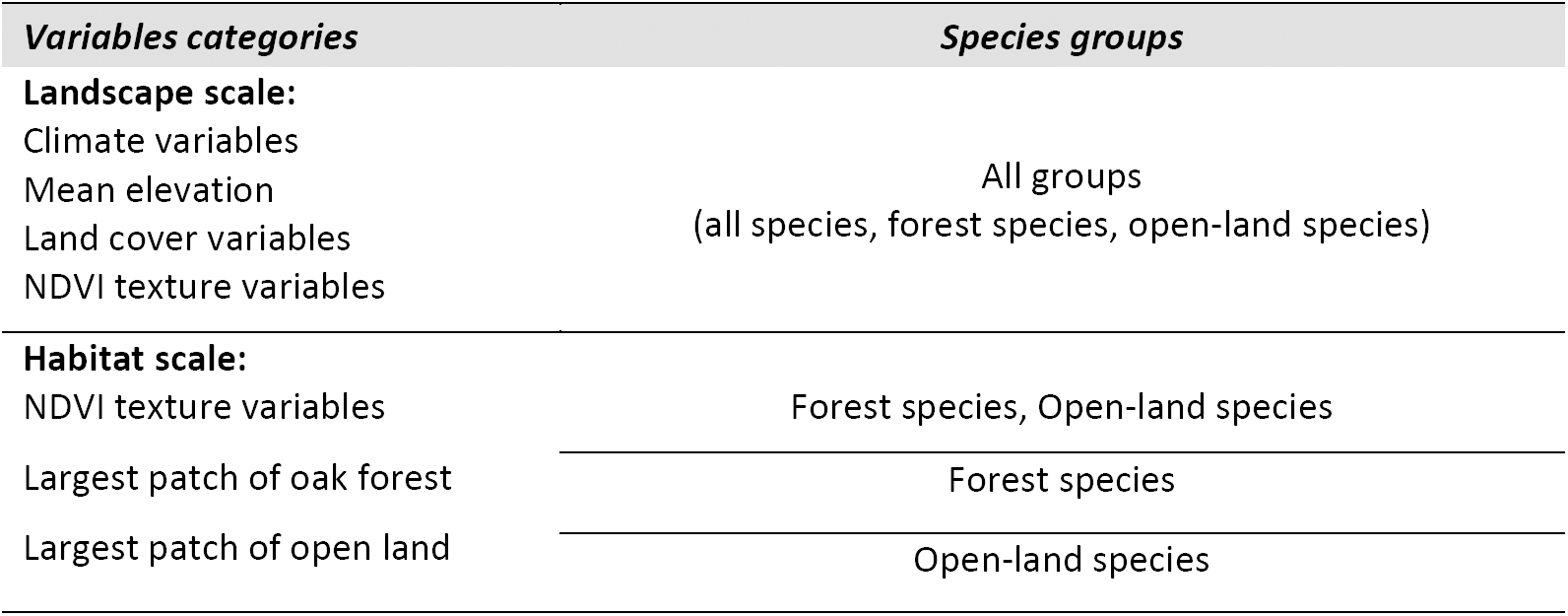
Main categories of candidate variables per species group. Landscape scale refers to variables measured for the full grid cell (i.e., using all pixels); habitat scale refers to variables measured using only the pixels overlapping patches of the preferred habitat, either forest or open-land habitats. Main habitats are the oak forest for forest species and open habitats for open-land species. NDVI texture variables were measured for the full cell (first-order variables) and using 3 x 3 and 9 x 9 pixel windows (second-order variables). Please see Appendix 2 for the full list of variables in each category.

### Climate, elevation and land cover data

Precipitation, temperature, and solar radiation data were collected from WorldClim database (http://www.worldclim.org/; Fick & Hijmans, 2017) on a 1 km resolution, elevation data were obtained from the *Digital Terrain Model* (30 m x 30 m) for Portugal in ArcGis *Online* 10.3.1 (https://www.arcgis.com/). Land cover data were extracted from the Portuguese land cover map, COS2007 (http://www.dgterritorio.pt/). For this study, we aggregated land cover classes into five categories: open land (permanent and temporary pastures, sand dunes, vineyards, shrubs and sparse vegetation), oak forest (open or closed forests and agro-forestry systems dominated by oaks), other forests (open and closed forests dominated by species other than oaks), urban (all the areas described at COS2007-level 1 as artificialized territory, including industries and roads), and water bodies (all the areas described at COS2007-level 1 as water bodies). To better capture species response to oak forest systems and reduce the influence of other forest types in the landscape, we defined eligible cells as those with a maximum of 20% cover of other forest types and where the cover of oak forest was not smaller than the cover of other forests (i.e., max 20% cover of other forest and oak forest cover ≥ other forest cover). The final sample included 59 cells.

### Remote sensing data

Six Landsat-5 images, free of clouds, corresponding to the following dates, paths and rows: 20^th^ March 2011 and 20^th^ July 2011 for path 203 and rows 33 and 34; 25^th^ April 2010 and 18^th^ August 2011 for path 204 and row 33, were used. These images were downloaded from the United States Geological Survey (USGS) Earth Explorer server, selecting the Landsat Surface Reflectance Level-2 Science Products (https://landsat.usgs.gov/landsat-surface-reflectance-data-products). Surface Reflectance products (at 30-meter spatial resolution) provide an estimate of the surface spectral reflectance as it would be measured at ground level in the absence of atmospheric scattering or absorption (Masek et al., 2006). After the availability of surface reflectance for all the images, the next step was the application of a cloud cover filter, obtained from visual inspection, to detect and mask clouds, cloud shades and haze. Two main RGB combinations were used: TM321 and TM453.

In order to monitor vegetation phenology, the Normalized Difference Vegetation Index (NDVI) was computed. This index is based on the normalized ratio between absorbed red light and near reflected infrared light (Rouse et al., 1973). NDVI values range from −1 (non-photosynthetically active vegetation) to +1 (highly photosynthetically active vegetation). This index has been used successfully in several studies to evaluate land cover performance (Li et al., 2004) or phenological information (Lloyd, 1990) or plant-community degradation (Alados et al., 2011) or to supply information about crops (Thenkabail et al., 1994; Martínez & Calera, 2001; Lyon et al., 2003; Jackson et al., 2004; Serra & Pons, 2008). Urban and water areas (NDVI ≤ 0) were masked from the original NDVI images to remove their signal from image texture analysis.

Image texture measurements quantify the spatial variation and arrangement of the reflectance values of neighboring pixels, expressing the level of spectral heterogeneity in a given area (Hernández-Stefanoni et al., 2012). Three first-order texture variables (NDVI entropy, mean NDVI, NDVI standard deviation) and two second-order texture variables (NDVI entropy, NDVI variance) were calculated for each cell and for the main habitat subsets, as listed in Table 1 and in Appendix 2.

First-order texture variables do not consider pixel neighbor relationships and are measured using the original image values within a certain group of pixels (Hernández-Stefanoni et al., 2012), which in our case was the full cell. On the other hand, second-order variables consider the spatial relation between neighboring pixels within a moving window (St-Louis et al., 2009), using a gray-level co-occurrence matrix (GLCM), which contain the probabilities of co-occurrence of values for pairs of pixels (Haralick et al., 1973). The *GLCM R* package was used to extract second-order texture measures within a 3 × 3 and a 9 × 9 moving window in four directions (0°, 45°, 90°, 135°). The *Zonal Statistics Tool* from ArcGIS 10.3.1 was used to summarize the mean and standard deviation and obtain a single texture value for each cell.

### Bird data

Georeferenced bird occurrence data was retrieved from the digital database of GBIF (Global Biodiversity Information Facility, http://www.gbif.org). We searched for species with resident populations in the study area and that use the various countryside habitats in the landscape, namely open-land habitats and oak forest (Appendix 3). The taxonomic nomenclature and the species distribution were retrieved from Catry et al. (2010). We excluded from our list the exotic species, species with wide habitats and high mobility, such as eagles and hawks, species with nocturnal habits, such as owls and nightjars, and insectivorous aerial birds, such as swallows and swifts (Godinho & Rabaça, 2011).

For the selected species, we collected species occurrences dated from April to June, to cover the birds’ nesting and reproduction periods (Pereira et al., 2015), between 2005 and 2015. Only records with no geospatial issues and under the *CCO 1.0* use license were used. The retrieved bird occurrence dataset, for the total number of searched years (i.e., 2005-2015), consisted of 122110 records.

### Selection of well-surveyed cells

Following, an identification of the cells for which species occurrences provided adequate inventories (Pineda & Lobo, 2009; Calderón-Patrón et al., 2013) was performed. For this purpose, data were analysed for each individual year, and aggregated in 2-year and 3-year time windows to increase sample size and coverage. The following steps were repeated for all possible 1-year, 2-year and 3-year contiguous time windows to identify well-surveyed cells. First, cells with less than 20 observed species were excluded (Chao & Jost, 2012). Second, two complementary methods were applied to estimate total cell species richness: non-parametric estimators based on the number of rare species (Chao 2 and Jackknife 1) (Hortal et al., 2006), and the number of estimated species at the 95% upper confidence interval of the accumulation curve produced with the Mao Tau analytical function (Mao et al., 2005). The three estimates were obtained with EstimateS 9.1.0 (Colwell, 2013). Based on the assumption that a higher number of records in a grid cell represents a higher survey effort, the total number of occurrence records in each cell was used as a surrogate of sampling effort (Hortal & Lobo, 2005).

Inventory completeness was calculated by relating the maximum estimated species richness among the three estimators (i.e., Mao Tau, Chao 2 and Jackknife 1) and the observed richness, that is, observed/maximum estimate × 100. Only the cells with completeness equal to or greater than 75% were considered as well-surveyed (Pineda & Lobo, 2009). From a total of 1839 grid cells in the study area (Figure 1), 1060 cells had GBIF records for the selected bird species. The number of cells suited for analysis decreased sharply after assessing inventory completeness. The best results (i.e., higher number of well-surveyed cells) were found for the 3-year time windows: 94 cells in 2007-2009, 91 cells in 2010-2012 and 68 cells in 2013-2015. By intersecting the 59 cells with adequate environmental data (see *Climate, elevation and land cover data* section) with the cells with well-surveyed bird data, we were able to match a maximum of 41 cells for the 2010–2012 time window (Figure 1, Appendices 1 and 4). Finally, after fitting generalized linear models (see section Data analysis) an over influential cell that had a Cook’s distance larger than 1 was detected. This cell was removed from the sample, resulting in a final sample of 40 cells (8042 records) that were used in data analyses.

### Species groups

The information in Pereira et al. (2015) was used to divide the 78 species present in our sample of 40 cells into two species groups: 27 forest species (10 specialists and 17 generalists) and 51 open-land species (8 farmland specialists, 12 farmland generalists, 22 edge species and 9 species requiring special landscape elements associated to farmland).

### Data analysis

From each set of candidate variables per species group (Table 1), we selected a group of non-collinear variables to be used in the statistical models. First, we calculated the Spearman’s pairwise correlations (*ρ*) among all pairs of predictor variables, and among predictor variables and the response variables (i.e., the observed species richness of each species group and all species) (Appendix 5). The predictor variables that were weakly correlated with the response variable (−0.1 < *ρ* < 0.1) were removed (Paudel et al., 2017). For the remaining predictor variables, if two or more variables were strongly correlated (*ρ* > |0.7|), we only kept the one most correlated with the response variable (Dormann et al., 2013; Paudel et al., 2017). Finally, we checked that the variation inflation factors (VIF) of the remaining predictors was lower than 5. The final sets of non-collinear variables are listed in Table 2. We also tested for the presence of spatial autocorrelation in the response variables using *Moran’s I statistic*. None of the response variables was spatially auto-correlated.

**Table 2.** Final set of non-colinear predictor variables per species group. Variables are ordered by the absolute value of their correlation with the species groups’ richness (ρ-Spearman rank correlation coefficients). Variables full name is presented in Appendix 2.

| *All Species* | $\rho$ | *Forest Species* | $\rho$ | *Open-land Species* | $\rho$ |
| --- | --- | --- | --- | --- | --- |
| Open areas (%) | -0.14 | Water areas (%) | -0.15 | Trimestral mean temperature | 0.10 |
| NDVI_sd_SP | 0.14 | NDVI_mn_SP | -0.17 | NDVI_var9x9_sd_OP_SP | -0.13 |
| Mean elevation | -0.20 | NDVI_ent9x9_sd_OF_SP | 0.20 | NDVI_ent9x9_mn_OP_SP | -0.14 |
| NDVI_ent9x9_mn_SP | 0.20 | NDVI_var9x9_sd_SP | -0.21 | NDVI_var9x9_mn_OP_SP | -0.15 |
| NDVI_mn_SP | -0.25 | Other forests (%) | 0.32 | NDVI_var3x3_mn_OP_SU | 0.17 |
| Other forests (%) | 0.26 | NDVI_var3x3_sd_OF_SP | -0.35 | Water areas (%) | 0.22 |
| Mean annual precipitation | 0.28 | NDVI_ent3x3_sd_SP | -0.37 | NDVI_mn_SP | -0.24 |
| Radiation Range | 0.30 | Urban areas (%) | 0.37 | NDVI_sd_SP | 0.25 |
| NDVI_var3x3_mn_SU | 0.36 | Radiation Range | 0.40 | Urban areas (%) | 0.32 |
| Max. temperature in June | -0.38 | NDVI_ent3x3_mn_OF_SU | 0.53 | NDVI_ent3x3_sd_SU | 0.35 |
| NDVI_sd_SU | 0.38 | Largest Patch_OF | 0.55 |  |  |
| Urban areas (%) | 0.48 | Min. temperature in April | 0.59 |  |  |
|  |  | NDVI_mn_SU | 0.82 |  |  |

Legend for the NDVI texture variables: mn – mean; sd – standard deviation; ent – entropy; var – variance; 3x3/9x9 – size of moving window; OF – Oak forest; OP – Open-land; SU – summer; SP - spring

Generalized linear models (GLM), with *Poisson error distribution* and *log link* function, were used to assess the importance of the environmental predictors in shaping species richness patterns. The variables listed in Table 2 were also tested for quadratic relationships. For each predictor variable, we compared the corrected Akaike Information Criterion (AICc) of the regression model comprising only the linear term of the predictor against the response variable, or the linear plus the quadratic term. If the regression model with the quadratic term had a better fit, both the linear and the quadratic term were included in the full GLM with all predictor variables. Quadratic terms were retained for mean elevation for all species and for percentage of water areas for the forest species group.

We used the *glmulti* package in *R* (Calcagno & Mazancourt, 2010) to test all possible combinations of the terms in the candidate set (i.e., full model) and rank the best models using AICc. Best models (difference from AICc _*minimum*_ < 2) were used to identify the most important variables affecting species richness patterns. Because we had a small sample size of 40 cells we only tested for main effects and restrained the maximum number of terms to be included in the candidate models to four. If a quadratic term was retained in one of the best models, the linear term was forced in the model (Vicente et al., 2013). We selected the most parsimonious model with the lowest AICc to check for overdispersion (*dispersiontest* function, in *R-package AER*); no evidence of overdispersion was found.

To identify the variables driving species turnover in bird communities, we applied generalized dissimilarity modelling using the *R-package gdm* (Manion et al., 2015). First, to fit GDMs, we used the final set of non-collinear variables to create a GDM site-pair matrix for each species group. In addition to the environmental predictors, geographical coordinates of cell centroids were included in the matrix to compute geographical distance. Variable significance was tested by combining Monte Carlo sampling and stepwise backward elimination as executed in the *gdm.varImp* function, with 250 permutations per step until only significant (α < 0.05) variables remained in the model. Second, to assess the amount of variance explained by each variable and the model, we fitted GDMs using *gdm* function, using only the significant predictors obtained with the *gdm.varImp* function. We used the default of three *I-spline* basis functions per predictor. We summed the coefficients of the *I-splines*, which are partial regression fits, to assess the relative importance of each predictor in determining patterns of beta diversity (Fitzpatrick et al., 2013). We also plotted the *I-splines* to visualize the variation and magnitude of species turnover along gradients of the significant predictors. The maximum height obtained by the curve represents the total amount of compositional turnover associated with that variable, holding all other variables constant. The slope of the I-spline indicates the rate of species turnover (Fitzpatrick et al., 2013).

To further test the explanatory role of the environmental variables found to have a significant effect on compositional dissimilarity, a correspondence analysis was applied to the matrix of species presence per site (i.e., cell), and the scores of the sites in the first principal axis were correlated against the environmental variables to test their association with species communities. Then, species scores were used to identify the species more related to the communities at the ends of the environmental gradient (i.e., with extreme scores).

## Results

### Species richness patterns

Best models for forest species richness suggest that mean NDVI in summer (NDVI_mn_SU), and radiation range are most important predictors of forest species richness at the landscape scale (Figure 2a, Table 3). Both variables are positively correlated with species richness (Table 2). For open-land species, the most important predictors were the standard deviation of second-order NDVI variance in 9×9 pixel windows in open land in spring (NDVIvar9×9_sd_OP_SP), the standard deviation of second-order NDVI entropy in 3×3 pixel windows in summer (NDVI_ent3×3_sd_SU), and the mean NDVI in spring (NDVI_mn_SP) (Figure 2b, Table 3). Spring variables are negatively correlated with species richness, while the summer variable is positively correlated (Table 2). Deviance explained by the best models varied between 66% and 67% for the forest species and was of 46% and 42% for the open-land species (Table 3). Total species richness was weakly described by the best models, with deviance explained varying between 33% and 30%, and no clearly dominant variables (Figure 2c, Table 3). Nevertheless, the mean of second-order NDVI variance in 3×3 windows in summer (NDVI_var3×3_mn_SU) and radiation range were the most important variables, both with positive coefficients.

**Table 3.** GLMs of species richness per species group, only the best models (ΔAICc ≤ 2) are shown. AICc and deviance explained (D^2^) provide a measure of models fit to data, with lower values of AICc and higher values of D^2^ indicating better fit. Models are ordered by the AICc. The linear term of mean Elevation (Elev_mn) was forced in the models for the All species group. The full name of the variables is presented in Appendix 2.

| | AICc | $D^2$ |
| --- | --- | --- |
| **Forest species** |  |  |
| SFor ~ 1 + NDVI_mn_SU + RadRg | 205.16 | 0.66 |
| SFor ~ 1 + NDVI_mn_SU + RadRg + NDVI_var9x9_sd_SP | 205.76 | 0.67 |
| SFor ~ 1 + NDVI_mn_SU + RadRg + NDVI_var3x3_sd_OF_SP | 205.93 | 0.67 |
| SFor ~ 1 + NDVI_mn_SU + RadRg + LgtPtch_Oak | 206.11 | 0.67 |
| **Open-land species** |  |  |
| SOpnL ~ 1 + NDVI_mn_SP + NDVI_var9x9_sd_OP_SP + NDVI_ent3x3_sd_SU + NDVI_ent9x9_mn_OP_SP | 220.45 | 0.46 |
| SOpnL ~ 1 + NDVI_mn_SP + NDVI_var9x9_sd_OP_SP + NDVI_ent3x3_sd_SU | 222.04 | 0.42 |
| **All species** |  |  |
| Stotal ~ 1 + Elev_mn + Elev_mn<sup>2</sup> + RadRg + NDVI_mn_SP + NDVI_var3x3_mn_SU | 267.99 | 0.33 |
| Stotal ~ 1 + Elev_mn + Elev_mn<sup>2</sup> + RadRg + NDVI_mn_SP + TmaxJ | 269.98 | 0.30 |

**Figure 2.**
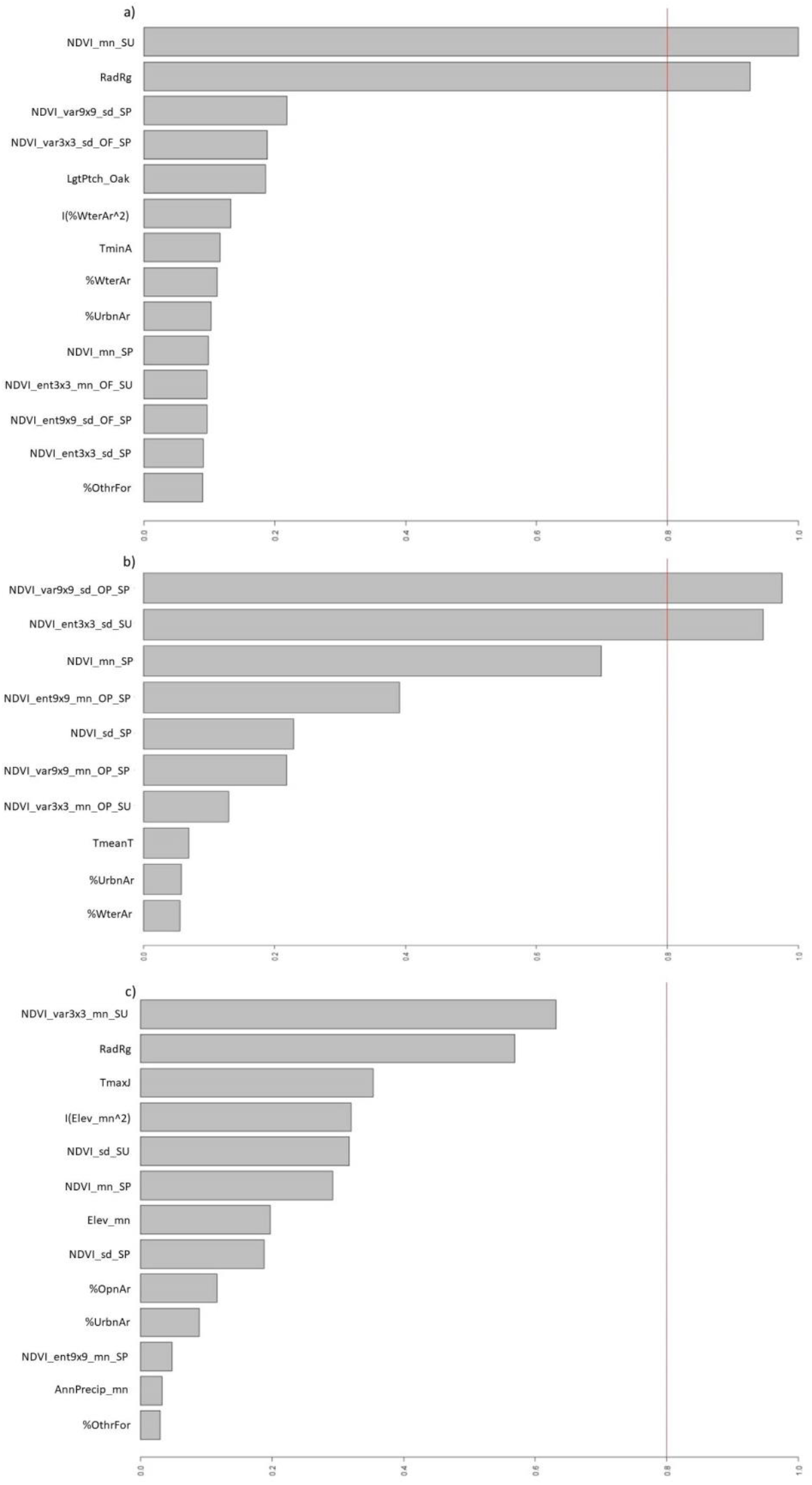
Average importance of model terms of generalized linear models. Plots show the overall support for each variable across all models. Model-averaged importance of predictors of a) forest species, b) open-land species and c) all species. Variables full name is presented in Appendix 2.

### Species dissimilarity patterns

The most important variables explaining species dissimilarity patterns at the regional scale (i.e., community dissimilarity between landscapes within the region) were mean NDVI in summer (NDVI_mn_SU) and geographical distance for forest species communities, and the standard deviation of second-order NDVI entropy in 3×3 pixel windows in summer (NDVI_ent3×3_sd_SU) and geographical distance for open-land species. For both species groups the compositional changes explained by the changes in vegetation structure, expressed by the gradient of the NDVI variable, mainly occurred below a threshold value becoming negligible above it (Figure 3 a,c). For instance, species turnover of forest species communities appears to stabilize above a threshold value for the NDVI_mn_SU, which is possibly related to the availability of arboreal habitat. Also, NDVI variables, related to vegetation structure, were more important than geographical distance in explaining compositional turnover, especially for forest species (Figure 3 b,d and Table 4). The GDM fitted for forest species explained 36.9% of deviance, while for open-land species the model explained 23.9% of deviance.

**Table 4.** Significant variables and their relative importance in the GDMs of forest species, open-land species and all species. Relative importance is determined by summing the coefficients of the *I-splines* from GDM. The percentage of null deviance explained by the fitted GDM model is also presented.

| *Significant variables* | *Relative importance* | *% Deviance explained* |
| --- | --- | --- |
| **Forest species** |  | **36.947** |
| NDVI_mn_SU | 0.514 |  |
| Geographic distance | 0.090 |  |
| **Open-land species** |  | **23.874** |
| NDVI_ent3x3_sd_SU | 0.240 |  |
| Geographic distance | 0.154 |  |
| **All species** |  | **39.844** |
| % OpnAr | 0.208 |  |
| NDVI_var3x3_mn_SU | 0.169 |  |
| Geographic distance | 0.122 |  |
| NDVI_sd_SU | 0.120 |  |

**Figure 3.**
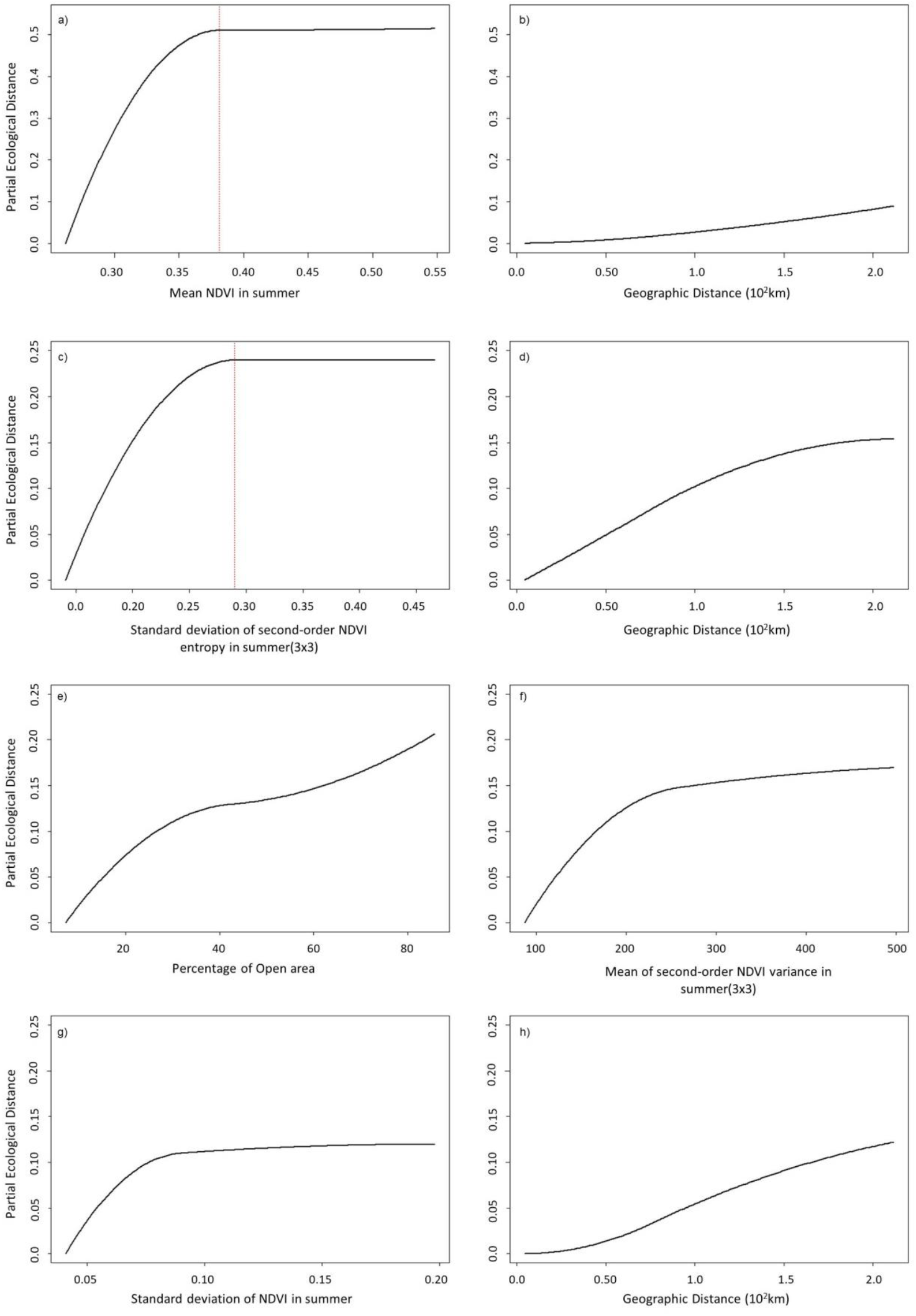
Generalized dissimilarity model-fitted *I-splines* (partial regression fits) for variables significantly associated with bird species turnover of forest species (a,b), open-land species (c,d) groups and total species (e,f,g,h). The maximum height reached by each function indicates the total amount of compositional turnover associated with that variable, holding all other variables constant.

The GDM model for all species (39.8% deviance explained) included a land cover variable (% cover of open land), a first-order texture variable (standard deviation of NDVI in summer), and second-order texture variable (mean of second-order NDVI variance in 3×3 pixel windows in summer, NDVI_var3×3_mn_SU) and geographical distance, all of similar importance (Figure 3 e-h, Table 4).

Site scores in the first principal axis (CA1) of the correspondence analyses conducted for forest species and open-land species were correlated with mean NDVI in summer (*ρ* = 0.82) for forest species, and with the standard deviation of second-order NDVI entropy in 3×3 pixel windows in summer (*ρ* = −0.64) for open-land species (Figure 4). Forest specialist species, such as *Phoenicurus phoenicurus, Aegithalos caudatus* or *Dendrocopus minor*, had high CA1 scores thus being more related to landscapes with high NDVI summer values (Figure 5a). For open-land species, the analysis reveals two groups: farmland species, such as *Tetrax tetrax* more associated to areas of low spatial heterogeneity (i.e., low values of NDVI_ent3×3_sd_SU) and edge species, such as *Emberiza cirlus* and *Petronia petronia*, more associated to areas of high spatial heterogeneity (Figure 5b). This analysis only included common species observed in at least 5 sites, which excluded three of the 27 forest species and 20 of the 51 open-land species.

**Figure 4.**
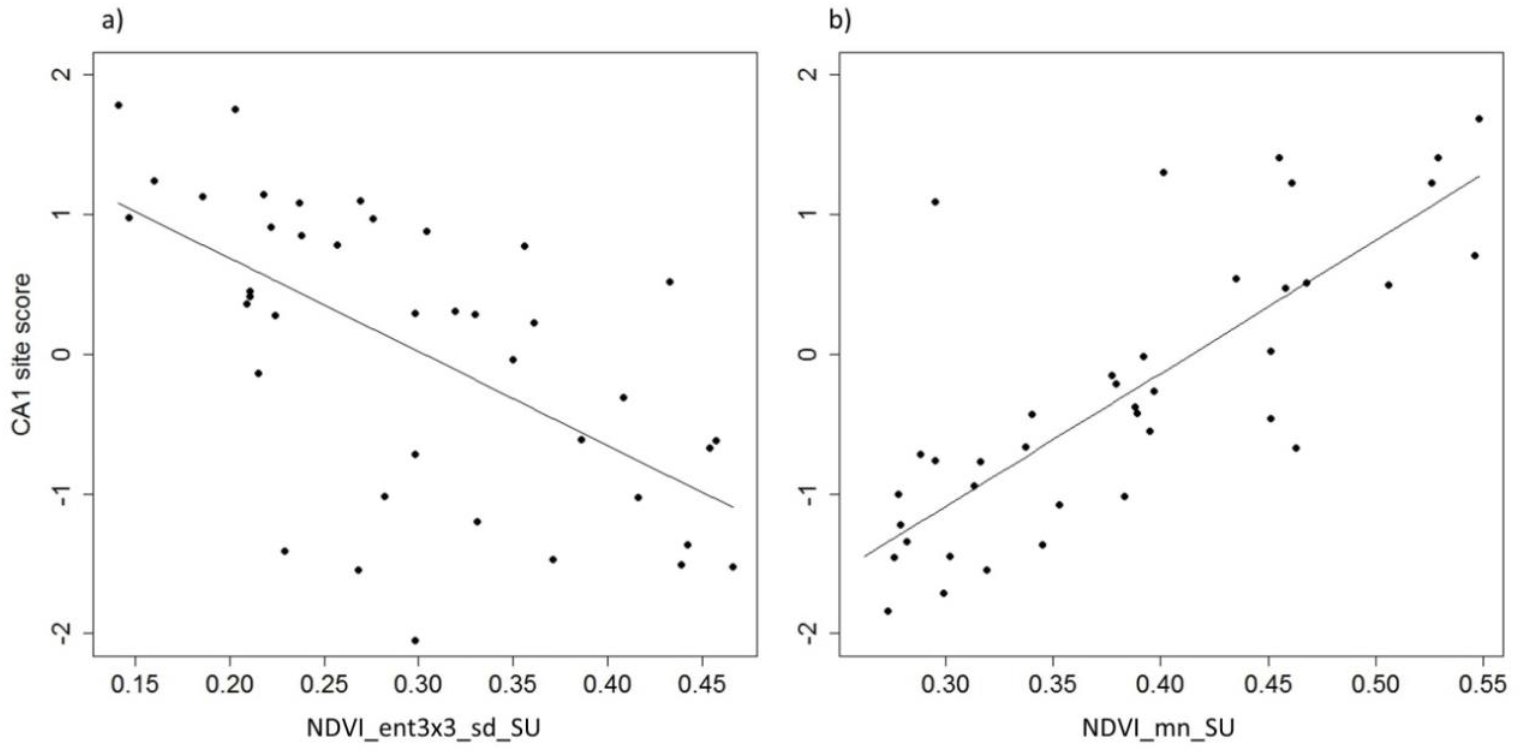
Relationship between site scores in the first principal axis (CA1) of the correspondence analysis and the environmental variables retained by the GDMs for forest species (a), *ρ* = 0.82, and open-land species (b), *ρ* = −0.64.

**Figure 5.**
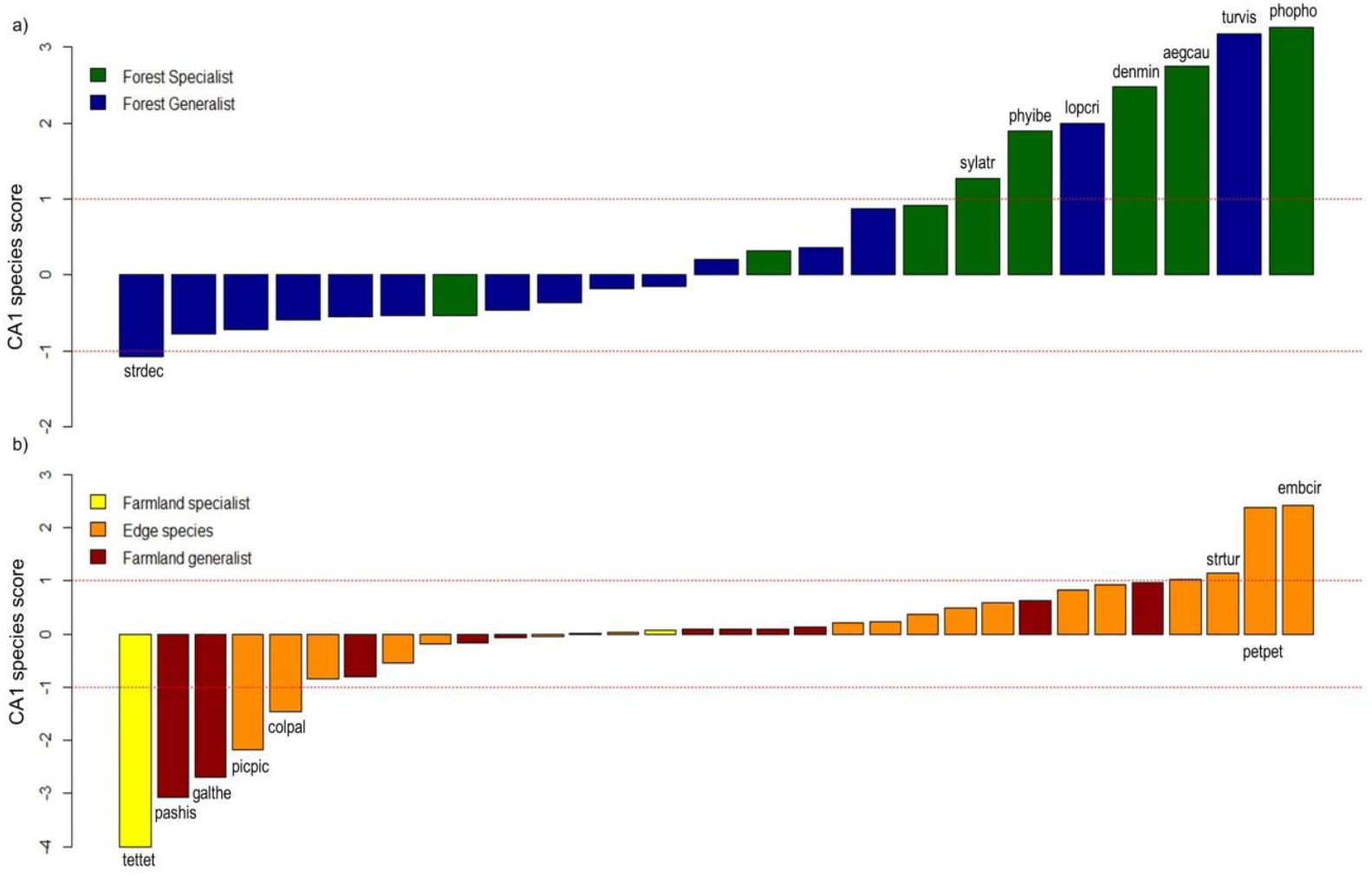
Species scores (a,b) in the first principal axis (CA1) of the correspondence analysis. Species with a score > 1 are identified (see Appendix 3 for the species full name).

## Discussion

Forest species richness was best described by models that included the mean NDVI in summer (NDVI_mn_SU) and the range of solar radiation in the landscape (here delimited by the grid cell). In Mediterranean systems, NDVI_mn_SU mostly captures the vegetation that keeps green in the dry summer season, that is, perennial trees and shrubs (Kuemmerle et al., 2006). NDVI_mn_SU was strongly correlated to forest species richness (rho = 0.82), but only moderately correlated to the proportion of oak forest in the landscape (rho = 0.56) and to the area of the largest patch of oak forest (rho = 0.60). While these land cover variables were only moderately correlated to richness (rho = 0.53 and rho = 0.55, respectively). Hence, NDVI_mn_SU probably provides a more accurate measure of the tree cover in the landscape, and therefore of habitat availability for forest species, being a better predictor of forest species richness. Wood et al., 2012 show that the mean of the NDVI is a good indicator of vegetation structure among habitats, because it captures the transitions in the landscape, especially if the habitats clearly differ in terms of structure, such as in grassland – woodland mosaics. The strong association between species richness and the mean of the summer NDVI could also be related to ecosystem productivity, as areas of high plant productivity would be associated to a higher availability of foraging resources (St-Louis et al., 2009). However, even if this is the case in our case study, the results are not explicit enough to confirm it. First, we did not find a similar association for open-land species or for the total species. Second, the correlation of species richness with the mean NDVI in spring (NDVI_mn_SP), which represents vegetation productivity at its peak, was not only weak but also negative for all the species groups, possibly because this variable is capturing the signal of both forest and open-land habitats. Regarding solar radiation range, this variable is related to the complexity of landscape morphology, characterized by parameters such as slope, aspect, and surface roughness. Hence, our results suggest that forest species, and their habitats, are more associated to landscapes with a more complex morphology, with diverse slopes and solar exposition that create the microclimate favorable to the growth of oak trees (Príncipe et al., 2014; Correia et al., 2015).

Contrary to our expectation forest species richness did not show a strong response to the structure of the main habitat but only to its availability in the landscape. This finding suggests that the availability of tree cover in the landscape may exert a stronger influence on species richness than the within-habitat structure of oak forest patches (e.g., tree density, tree cover arrangement). A similar result, also for Alentejo, was reported by Santana et al., 2017, the authors found a strong association of woodland bird species richness to the amount of woodland in the landscape but not to landscape configuration. This finding is further supported by the GDM results, which suggest that the composition of forest communities tends to stabilize, with limited addition of new species or species replacement, above a threshold value of the mean of NDVI in summer (Figure 3 a). On the other hand, our results do not reveal a significant response of forest birds to within-habitat structure, previous studies have reported such effect. For instance, grazing management and changes in tree density have been reported to affect bird communities in wood-pastures (Hartel et al., 2014; Godinho, 2016). The lack of response revealed by our analysis may be related to the resolution of the data used. Bird data were aggregated for the landscape mosaic and the optical satellite data probably misses most or part of the signal of the forest understory, even if understory features are correlated to canopy features or affect the reflectance of the canopy layer (Culbert et al., 2012; Wood et al., 2012). Moreover, the selection of the NDVI_mn_SU by both the GLM and the GDM analyses, suggests that the fitted GDM is capturing better the compositional changes related to changes in species richness, rather than the changes related to species replacement. This is possibly associated to the limitations of the data used as described above, but also to the extent of the study area. The Alentejo region may not be wide enough to observe species replacement when communities within landscapes with identical forest cover are compared. In fact, the effect of geographic distance on species replacement was found to be smaller than the effect of landscape structure for forest species (Figure 3 b). On the other hand, the species found to be more associated to high NDVI_mn_SU values were forest specialists (Figure 5 a), such as *Phoenicurus phoenicurus, Aegithalos caudatus* and *Dendrocopos minor*, suggesting that these species preferentially occur in landscapes where NDVI_mn_SU, an indicator of tree cover, reaches higher values.

Open-land species richness was best described by models that included second-order texture variables related to the arrangement of the vegetation in the main habitat and in the landscape mosaic. Namely, species richness was negatively correlated with the standard deviation of the second-order NDVI variance (9 × 9 window) measured in open land in spring (NDVI_var9×9_sd_OP_SP) and positively correlated with the standard deviation of the second-order NDVI entropy (3 × 3 window) measured in the landscape in summer (NDVI_ent3×3_sd_SU). This suggests that, on the one hand, the richness of open-land species is promoted by vegetation homogeneity within open-land habitats (i.e., richness was higher when the moving windows of 9×9 pixels in the open-land patches shared similar levels of NDVI variance), and, on the other hand, by vegetation heterogeneity at the landscape scale (i.e., the richness of open-land species is higher in landscapes where the entropy of NDVI measured in moving windows presents high variation at the landscape scale, ranging from low to high entropy). High values of NDVI_ent3×3_sd_SU correspond to landscapes with sharp transitions between land cover patches, for instance where homogenous crop land is delimited by edge habitats, or where different types of crops, planted in patches, coexist in the landscape. This pattern agrees with the results reported by Santana et al., 2017, who also found a strong association between farmland species richness and edge density in Alentejo.

Other studies also report a significant correspondence between how bird species use the space and its resources and habitat structural features, such as matrix of small patches of different habitats (Godinho & Rabaça, 2011; Hartel et al., 2014; Pereira et al., 2015). Because the correlation between open-land species richness and NDVI_var9×9_sd_OP_SP was weak (rho = −0.13), the association between this variable and species richness should be interpreted with caution. Regarding NDVI_ent3×3_sd_SU, this variable was moderately correlated to open-land species richness (rho = 0.35) and was co-linear to the proportion of open-land habitats (rho = 0.81) and to the area of the largest patch of open-land (rho = 0.81), thus being a good indicator of open-land habitats. NDVI_ent3×3_sd_SU was also co-linear to many of the entropy variables tested in this study.

As observed for forest species, the turnover in open-land species communities was mostly explained by the gradient of a key predictor of species richness, here NDVI_ent3×3_sd_SU (Figure 3 c). Also, community composition appears to stabilize above a threshold value. Again, this pattern suggests that at this scale of analysis, and with the data available, the GDM is mostly capturing the compositional changes related to changes in species richness and, as discussed below, to different types of open-land habitats. Nevertheless, the deviance explained by the model and the relative importance of NDVI_ent3×3_sd_SU are relatively low (Table 4), leaving a large share of compositional dissimilarity unexplained. As found in previous studies (e.g., Reino et al., 2009; Stoate et al., 2009; Santana et al., 2017) the composition of open-land species communities is very responsive to management and to the composition of crops, factors that were not tested in this study but likely to be relevant. Another important aspect regards the structure of the species group, which encompasses both farmland and edge species that have different ecological preferences within the open-land habitats. This may prevent the detection of a strong response pattern for the whole group. In fact, farmland species appear to be more associated to landscapes with high values of NDVI_ent3×3_sd_SU (Figures 4 and 5b), which are also richer in species, and edge species to landscapes with low values of NDVI_ent3×3_sd_SU (Figures 4 and 5b). For example, in Alentejo, *Emberiza cirlus* prefers the edges or clearings of forested patches and the presence of understory shrubs seems to favour its occurrence (Catry et al., 2010).

Finally, because forest and open-land species respond differently to landscape attributes, the best models for total species richness were less effective in explaining richness patterns than the models for the species groups (Table 3). Overall, total species richness appears to be higher in landscapes with high heterogeneity at the local scale and the landscape scale, as described, respectively, by the NDVI_var3×3_mn_SU and the radiation range (Figure 2). This result agrees with previous studies that reported the effect of habitat heterogeneity in species richness (St-Louis et al., 2009; Leal et al., 2011; Martins et al., 2014). In contrast to species richness patterns, species dissimilarity was well described by the GDM for total species (Table 4). This is not surprising since the two species groups show distinct preferences in the use of the landscape, and therefore the patterns of species replacement are clearer and captured more straightforwardly by the data. The turnover in community composition is well explained by a land cover gradient between forest and open-land habitats (note that the two variables are inversely correlated in our sample), with species replacement slowing down (i.e., flatter slope) at intermediate levels of the gradient (Figure 3 e). NDVI_var3×3_mn_SU and NDVI_sd_SU were also identified as significant predictors of community dissimilarity. The effect of these texture variables is probably related to changes in species richness driven by landscape heterogeneity. Also, both are summer variables which highlights the importance of perennial vegetation in shaping landscape structure and providing habitat diversity for bird communities.

## Concluding remarks

In a time of fast environmental change, the assessment of biodiversity response at large spatial scales, fine spatial resolution and frequent time intervals is necessary but limited by operational costs (Nagendra et al., 2013; Pettorelli et al., 2016; Proença et al., 2017). The development of cost-effective and expedite methods to monitor biodiversity change is therefore required. In this study, we used open access satellite imagery and biodiversity data to characterize vegetation structure and bird communities in rural landscapes in the Alentejo region in Portugal. Our results show that bird communities respond to vegetation structure, measured using NDVI texture variables with a 30m-pixel resolution, and that these variables can predict habitat availability more accurately than variables of habitat extent and structure based on land cover maps. Moreover, in seasonal ecosystems, such as the Mediterranean ecosystems, the date of the satellite imagery is important since images collected at different seasons will capture the vegetation at different phenological stages and therefore the habitat elements that are critical for different species groups. We found that summer images, when the green perennial vegetation becomes more apparent, are particularly suited to model the diversity patterns of forest species, because distribution of tree cover in the landscape is well captured. Summer data is also useful to monitor landscape structure and the perennial elements that shape the habitat of open-land species. On the other hand, spring images appear to be relevant for open-land species to monitor vegetation within open-land habitats, but the adequacy of the data at the landscape scale is affected by the simultaneous reflectance, at peak productivity, of the herbaceous and the woody components. The extraction of the herbaceous component from spring images could provide a solution to work around this limitation (Kuemmerle et al., 2006). Regarding biodiversity data, the use of open-access stored at GBIF enabled the assemblage of large-scale dataset. However, because these data originate from diverse primary sources and were collected using different methods, the process of data filtering, validation, aggregation and assessment of data quality prior to data analysis is essential. We successfully developed a protocol to follow these steps and ensure data quality before analyzing species diversity patterns. Although a vast amount of bird occurrence data is available for the Alentejo, we still had to aggregate data spatially and temporally in order to assemble datasets with adequate inventory completeness and ensure spatial accuracy. The consequence of data aggregation was a reduction of the spatial resolution of the bird data, which, as discussed above, may have impaired the detection of species responses to vegetation structure at lower scales. On the other hand, we were able to successfully conduct a regional scale analysis only informed by open-access data, which illustrates the potential of these data to be used in large scale biodiversity monitoring.

Hence, the combination of open-access biodiversity data and satellite imagery can be used as a cost-effective option to monitor biodiversity response to changes in vegetation structure. In particular, our findings suggest that NDVI texture variables are good descriptors of vegetation structure and can be used to monitor habitat change and predict changes in species communities.

## Acknowledgements

This study was supported by FCT/MCTES (PIDDAC) through project UID/EEA/50009/2013, by the H2020 project ECOPOTENTIAL (Grant Agreement no. 641762), and by the ICNF (AD 288/2014/ICNF/SEDE). V.P. was supported by FCT (grant SFRH/BPD/80726/2011).

**Appendix 1.**
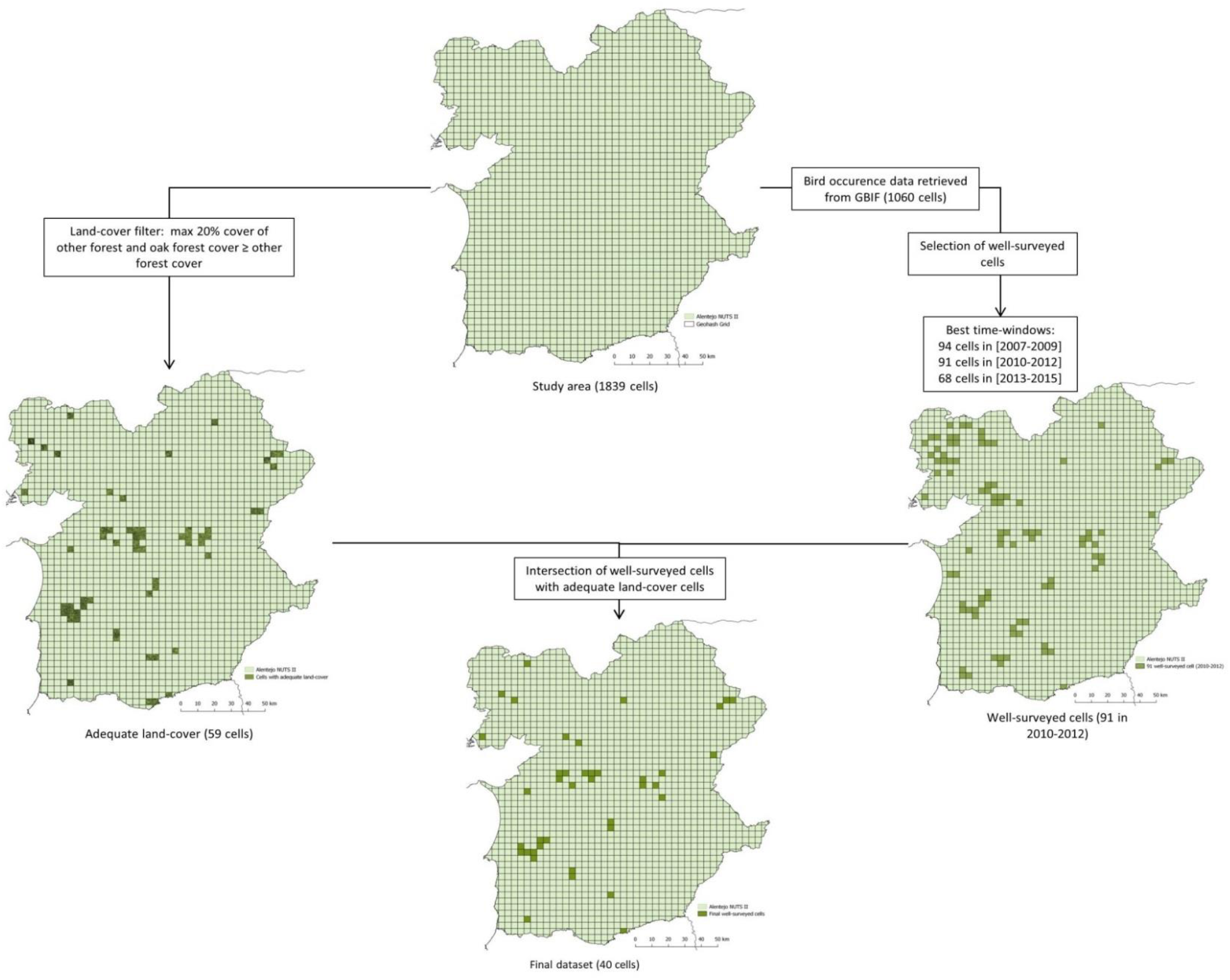
Summary of data filtering steps to obtain the final sample of 40 cells suitable for data analysis. The intersection of the cells with adequate land cover with the best time windows, resulted in a maximum number of 41 cells for the 2010-2012 time window. However, after fitting generalized linear models (see section Data analysis) an over influential cell that had a Cook’s distance larger than 1 was detected. This cell was removed from the sample, resulting in the final sample of 40 cells (8042 records).

**Appendix 2.**
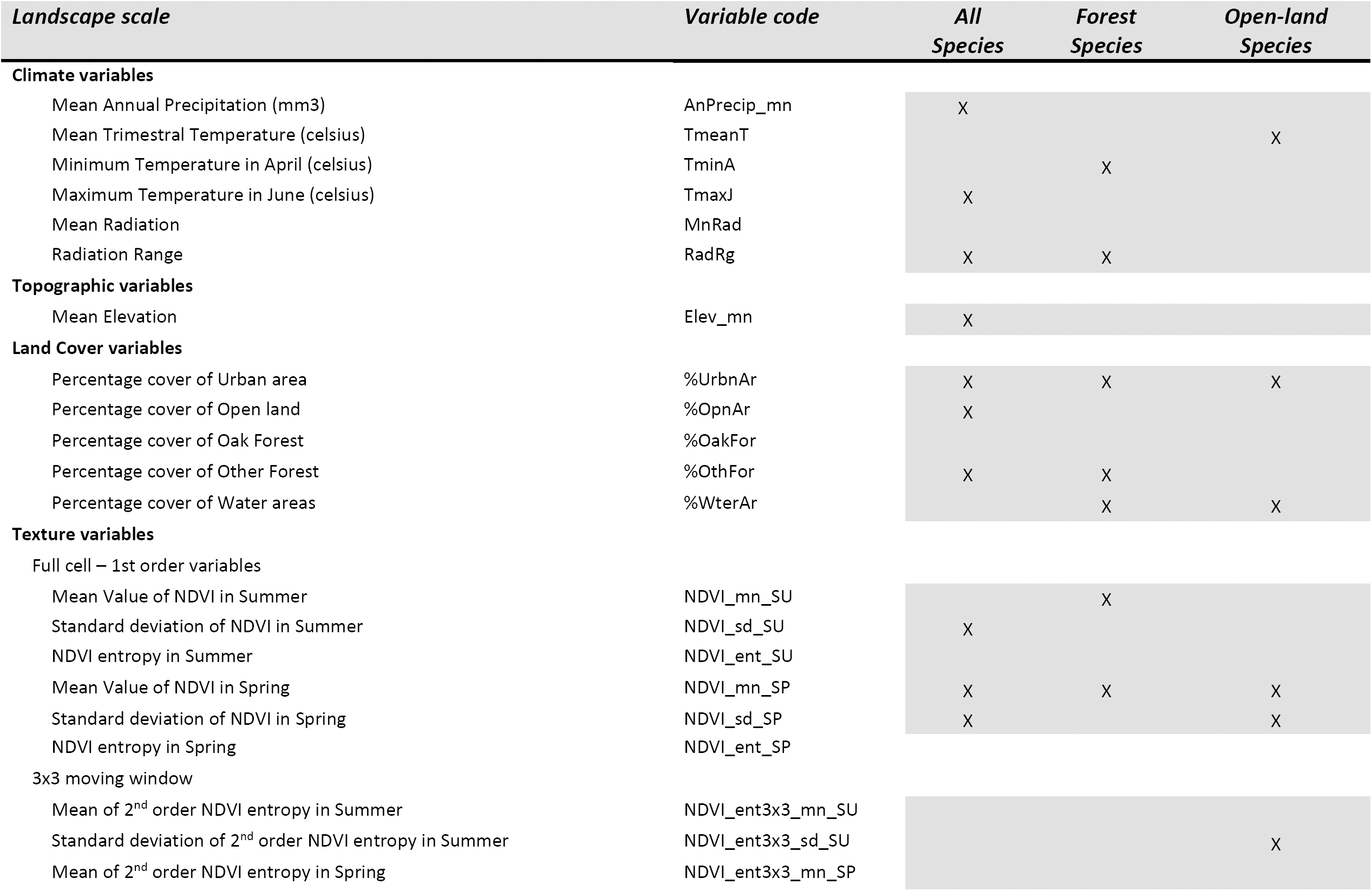
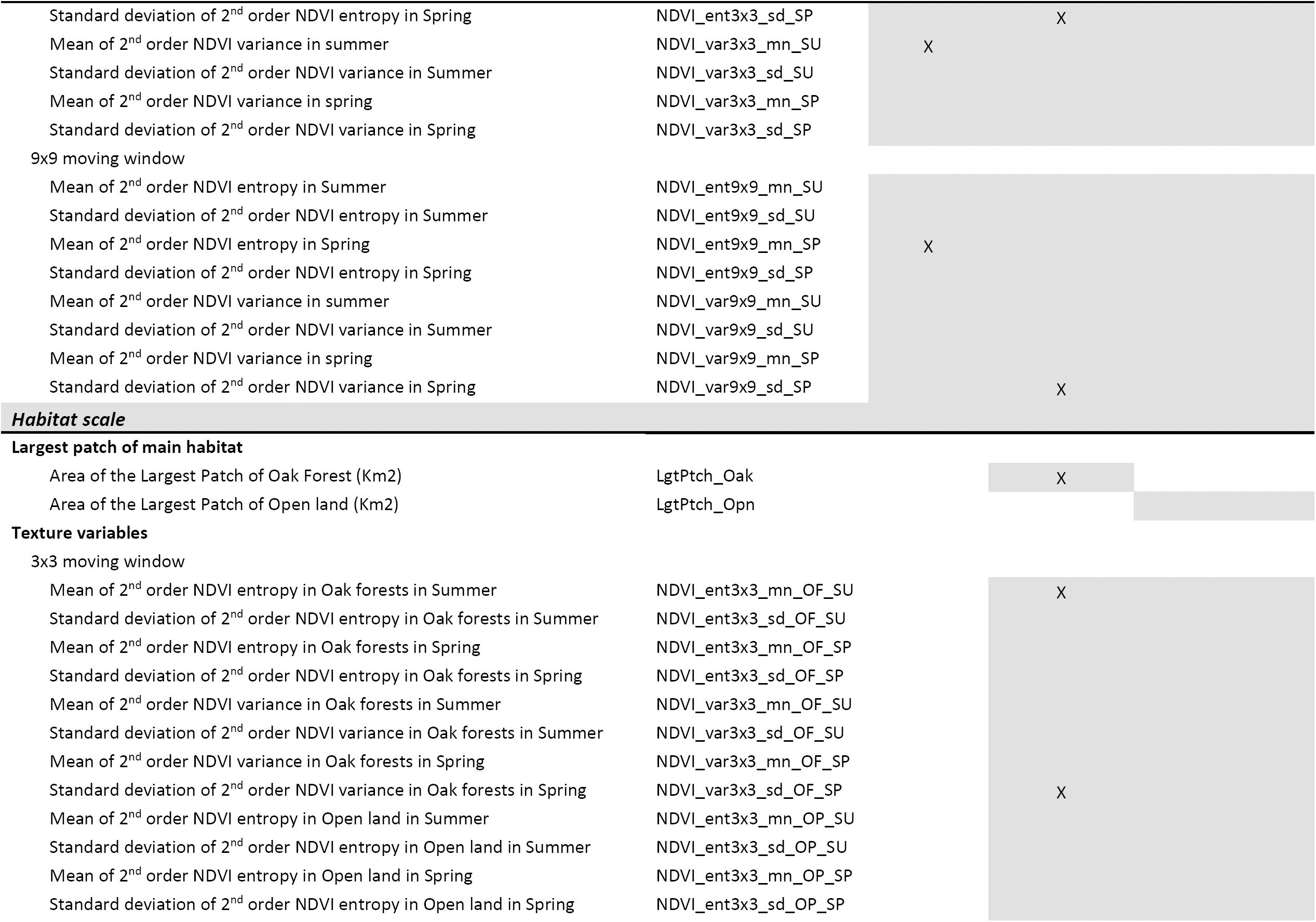
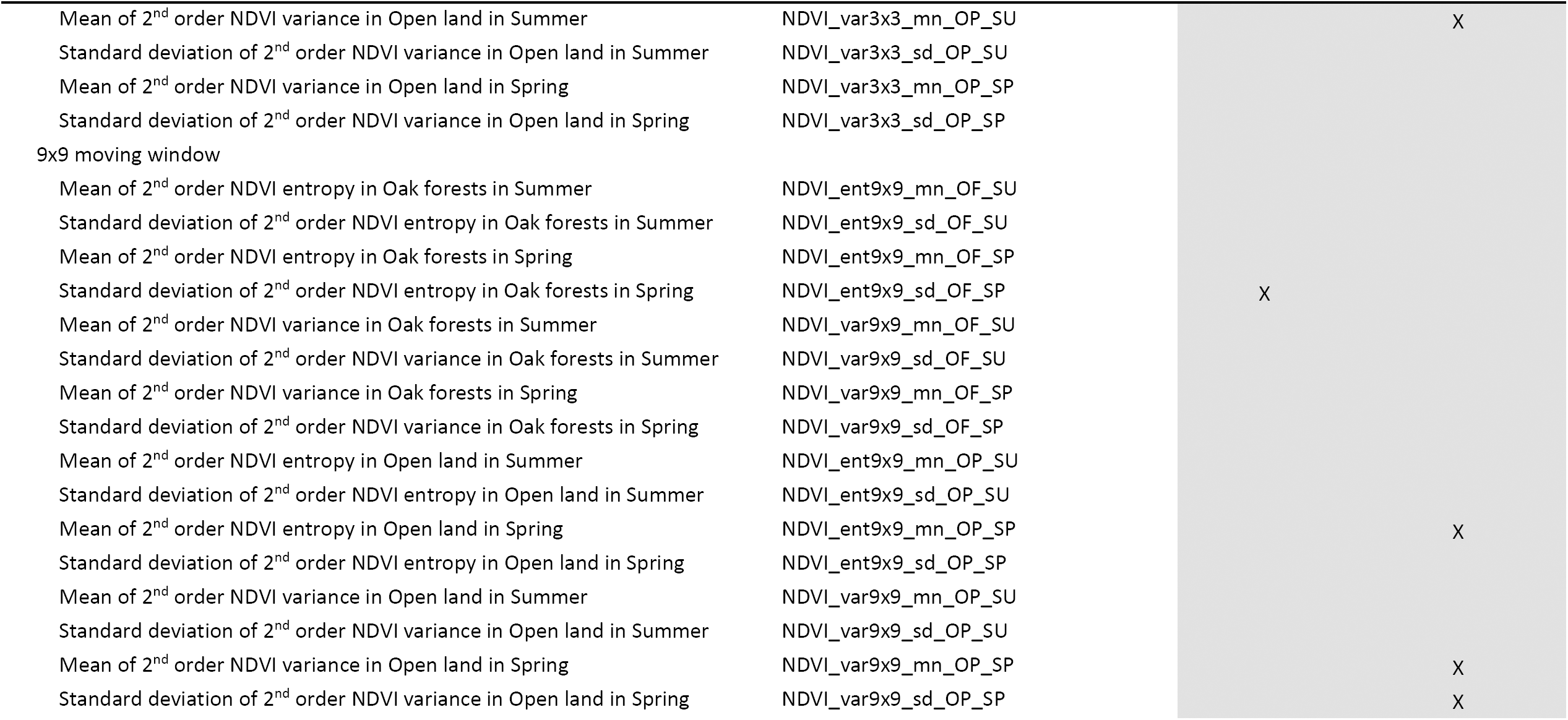
Full list of candidate variables (shaded cells) in each category per species group and the final list of non-colinear significant variables retained after variable selections (marked with “X”) per group.

**Appendix 3.**
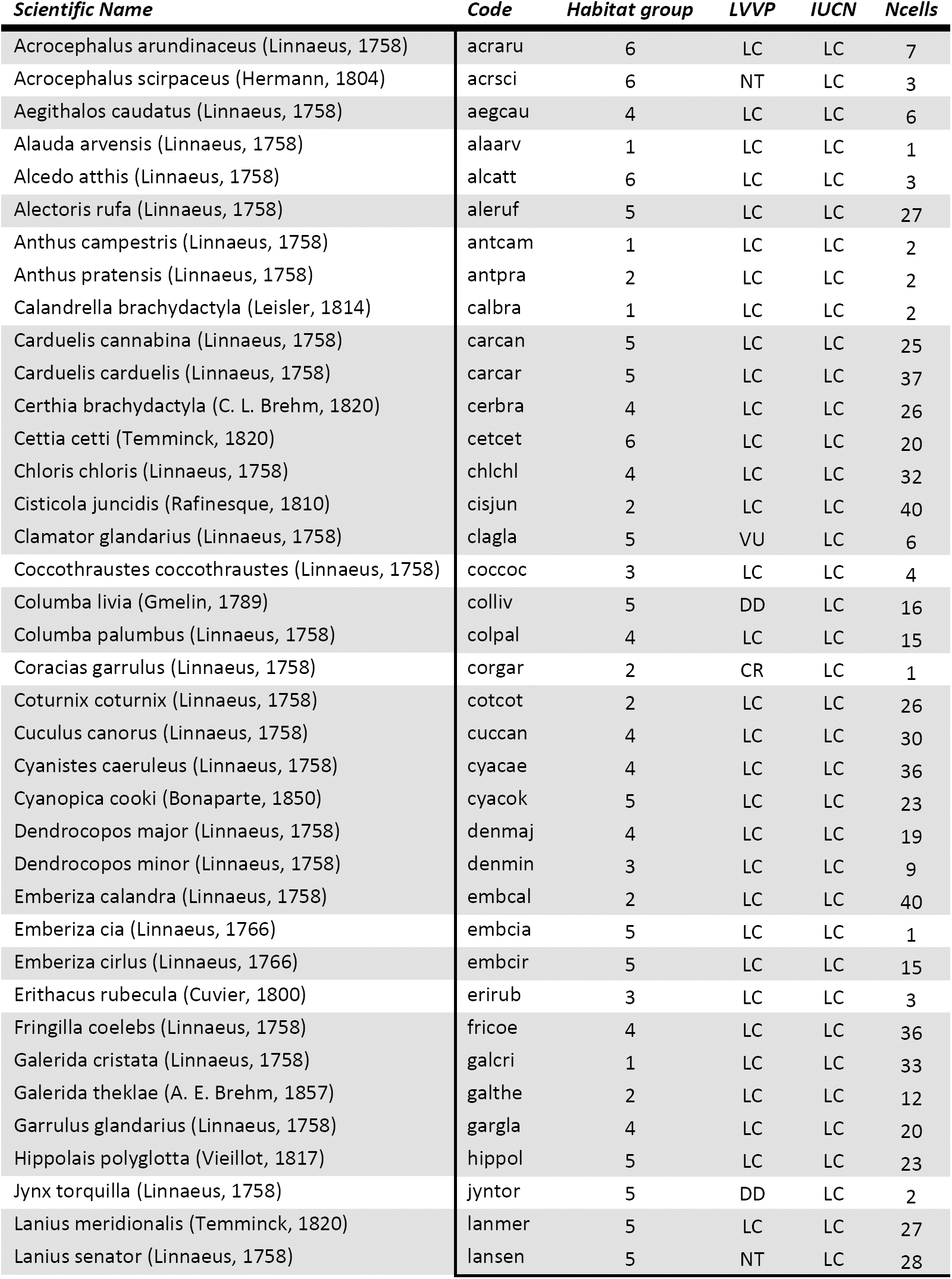
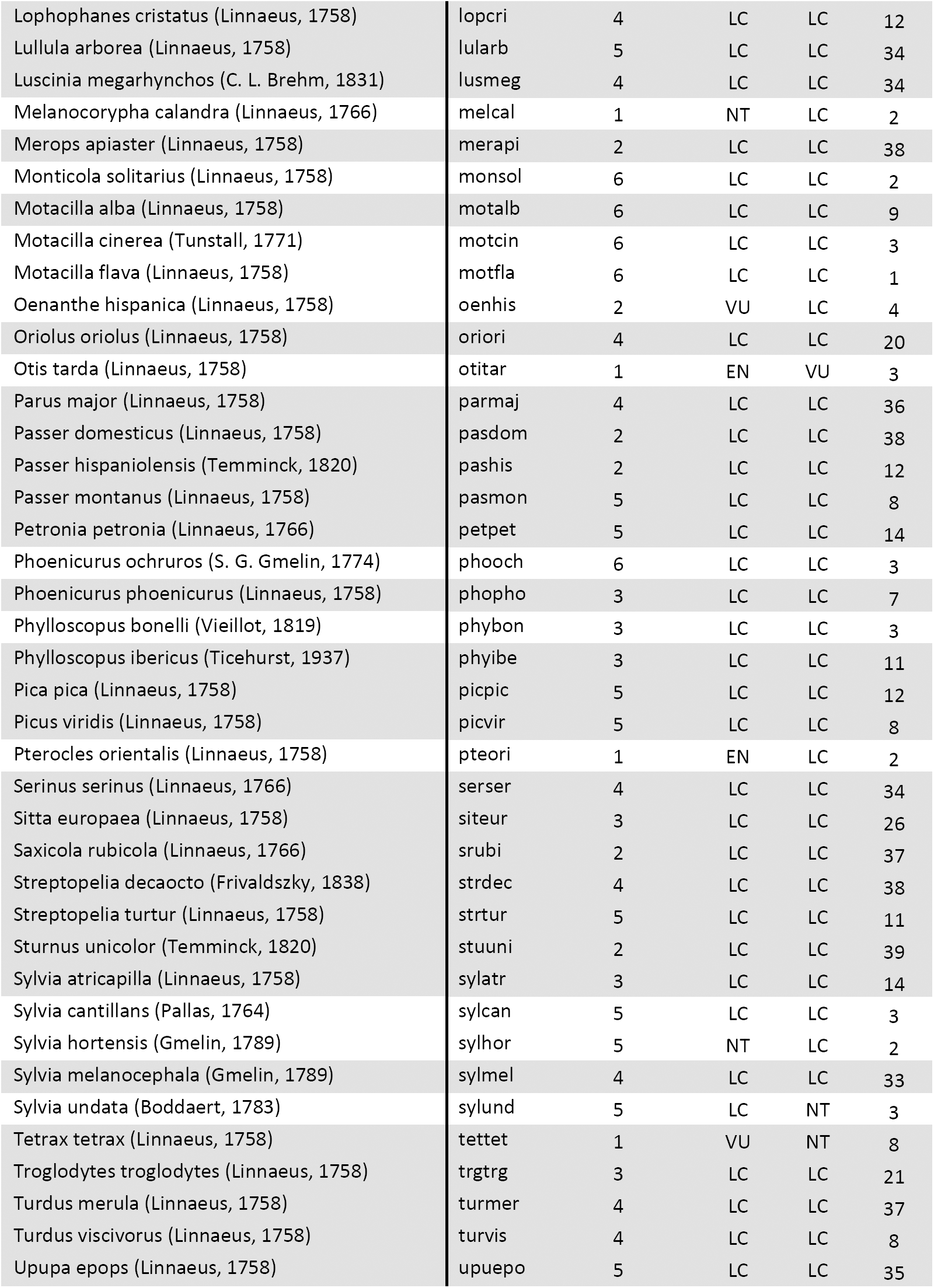
Full species list used in this study. Grey shadowed lines represent the species present in at least 5 cells, used in the correspondence analysis. The conservation status is also shown for the IUCN Red List and for the Portuguese Red List (LVVP). Habitat group: 1 – Farmland specialists, 2 – Farmland generalists, 3 – Forest specialists, 4 – Forest generalists, 5 – Edge species and 6 – Special elements.

**Appendix 4.**
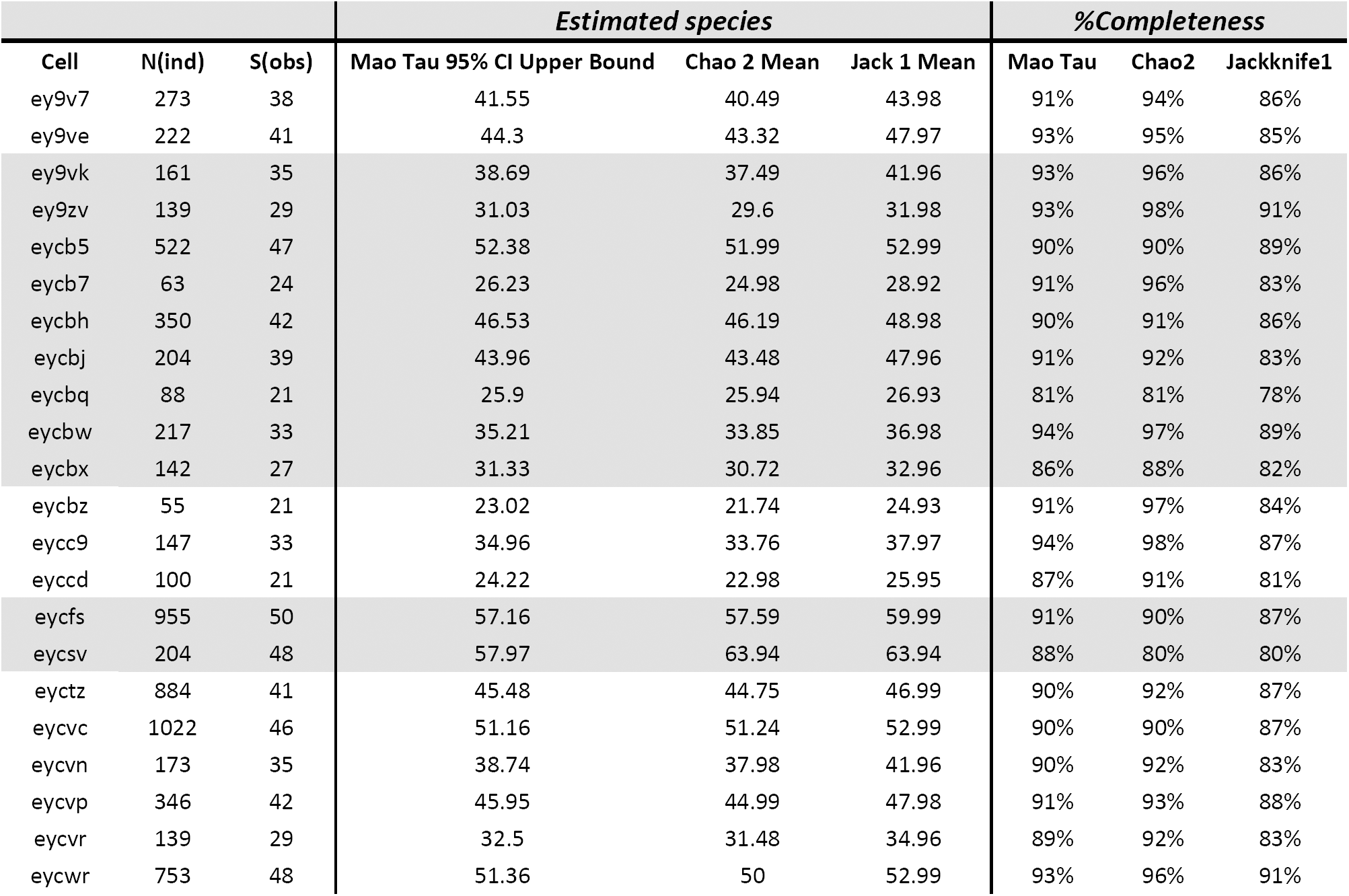
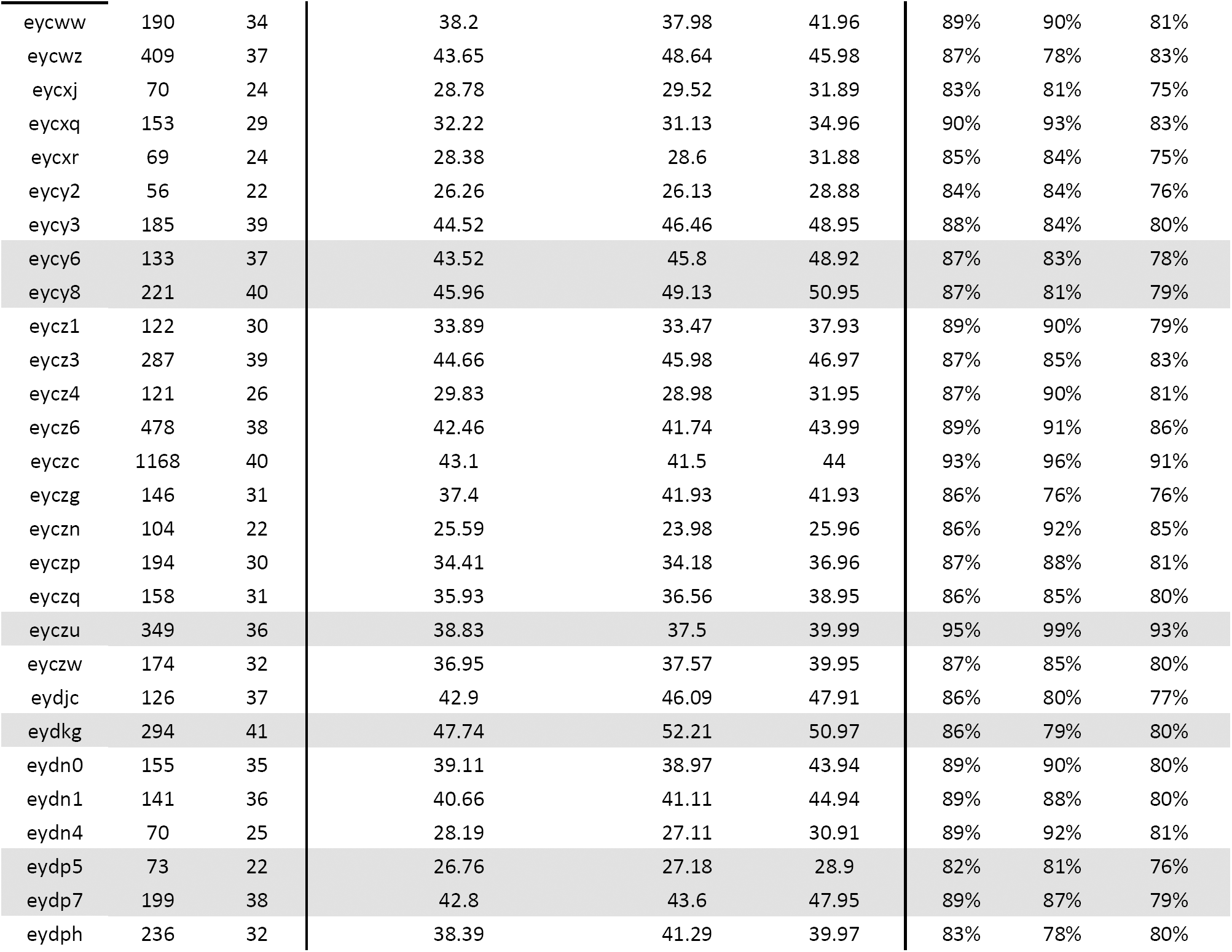
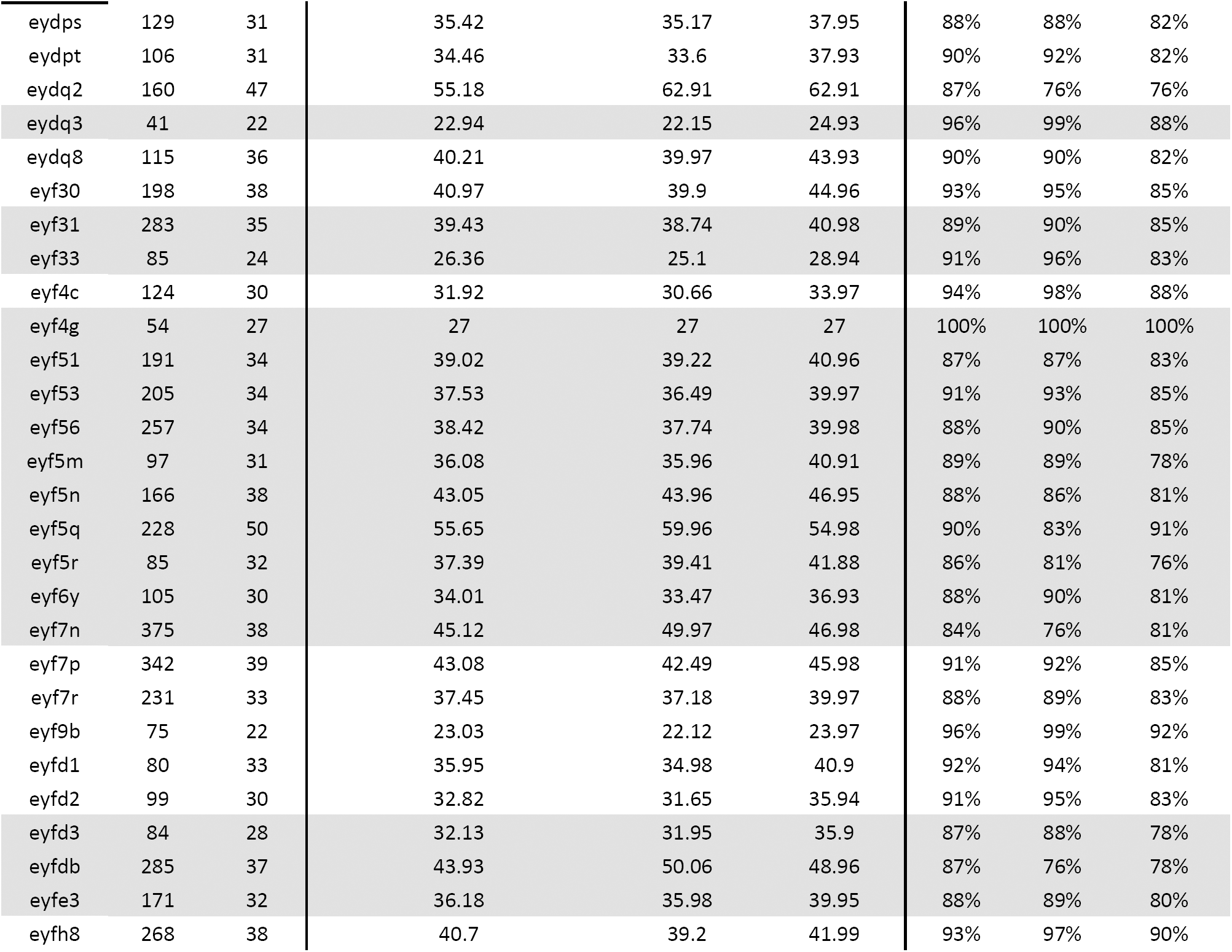
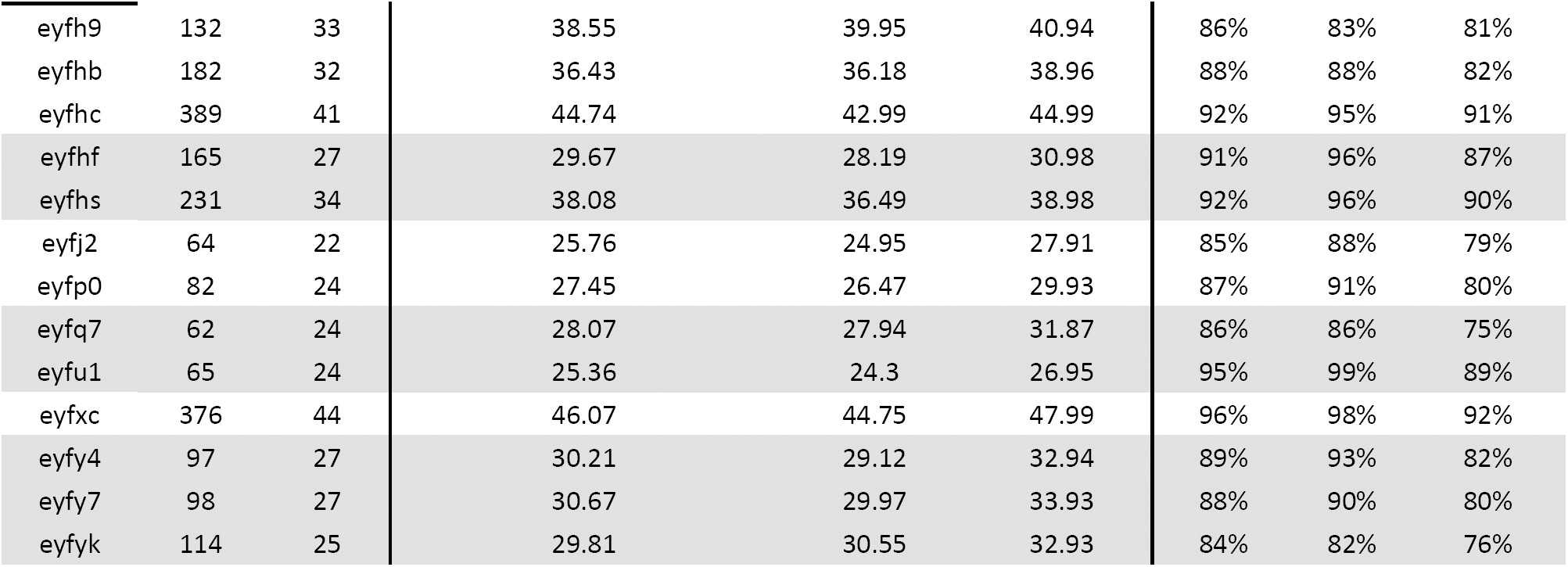
Total list of well-surveyed cells in the 2010-2012 time window (n= 91 cells). From this list, only 40 cells (shadowed in grey) were selected for data analysis after intersecting with the 59 cells with adequate land cover data.

**Appendix 5.**
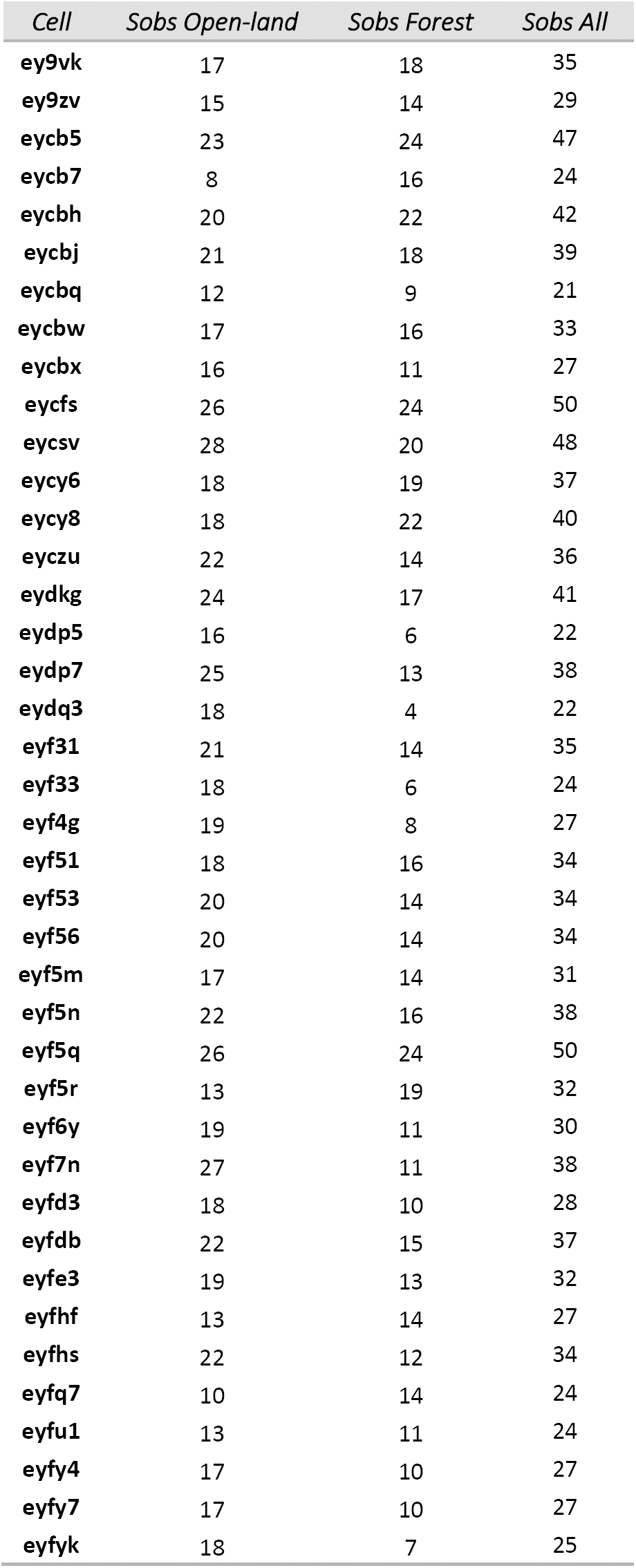
Observed species richness per species group and for all species in the final 40 selected cells.

